# Day and night posture of the bluegill sunfish (*Lepomis macrochirus*)

**DOI:** 10.1101/2023.07.13.548884

**Authors:** Michael Fath, Eric D. Tytell

**Affiliations:** Tufts University

## Abstract

Many animals assume characteristic postures when resting or sleeping. These postures are often stable and can be maintained passively, thus reducing the energy cost for maintaining an unstable posture. For example, many tetrapods lay prone on the ground and some negatively buoyant fishes are also able to rest on the substrate. Other fishes rest suspended in the water column. Counterintuitively, hovering this way can be of similar energetic cost to swimming. Even if the fish is perfectly neutrally buoyant, any displacement between its center of mass and center of buoyancy will produce destabilizing pitching torques that the fish must constantly work to counteract if they wish to maintain that posture. We hypothesized that a neutrally buoyant fish could rest at an equilibrium – a posture at which no destabilizing torques are produced by the body --to minimize the metabolic costs associated with hovering. Specifically, we studied the bluegill sunfish (*Lepomis macrochirus*), which is unstable in a horizontal posture. However, by pitching their bodies up or down they may be able to attain a less costly equilibrium posture, one which vertically aligns their center of mass and center of buoyancy. To test this hypothesis, we measured pitch angle of bluegill over the course of 24 hours. We also measured the pitch angles of the body that correspond to stable and unstable equilibria. We found that the stable equilibrium was a belly-up posture, and the unstable equilibrium is a dorsal side up posture pitched 53±26° head-down. The fish rested at a head-down pitch of -10.7±0.4° degrees, which is significantly steeper than the average pitch during the day of -3.4±0.8° degrees head down. These results show that bluegill do not rest at unstable or stable equilibrium. However, they do rest closer to unstable equilibrium at night than during the day. This may allow them to decrease destabilizing torques generated from the relative locations of the COM and COB while maintaining maneuverability.

## Introduction

The ability to move is one of the defining characteristics of animals. Movement allows animals to seek out food, shelter, and mates at geographically distant locations (Dorfman et al., 2022; Nathan et al., 2008; Pyke, 1983). However, movement comes with a metabolic cost. One way to reduce that metabolic cost – particularly when resources are scarce or predators are present—is by resting or sleeping (Schmidt, 2014; Siegel, 2011). Resting is associated with periods of immobility and passively maintained postures across many vertebrate and invertebrate taxa (Joiner, 2016; Keene & Duboue, 2018), thus reducing energy spent locomoting or maintaining a posture (Siegel, 2011). Fishes, both bony and cartilaginous, alter their behavior while resting, including by assuming stereotypic postures (Kelly et al., 2020). Negatively buoyant fishes can rest on the substrate (Kelly et al., 2022; Shapiro & Hepburn, 1976), neutrally buoyant fishes may assume a non-horizontal posture (Zhdanova et al., 2001), and some positively buoyant fishes can rest against the ceilings of underwater caves (Ciancio et al., 2016). Ciancio et al. (2016) proposed that changes in posture while resting are associated with reducing the energy required to maintain an unstable posture in fishes.

Fishes in a dorsal-side-up posture are often unstable in pitch (medio-lateral axis rotation) or roll (long axis rotation; Fath et al., 2023; Fish & Holzman, 2019; Webb & Weihs, 1994). This instability arises from a separation between the center of mass (COM), where the downward force of gravity is located, and the center of buoyancy (COB), where the upward force of buoyancy is located (Weihs, 1993). Any horizontal distance between the COM and COB generates torques which a fish must actively counter to maintain a horizontal, level posture (Eidietis et al., 2002; Webb, 2004). As discussed above, some fishes relax their posture while resting, but fishes which rest in the water column must actively maintain an unstable posture. Biomechanical studies of fish resting are rare, but some indicate differing kinematics between day swimming and night swimming (Goldshmid et al., 2004).

Hovering presents a potentially high metabolic cost for fishes which do not rest against a substrate. For birds, the metabolic rate during hovering is higher than flying at moderate speeds (Clark & Dudley, 2010). There is evidence that in some fishes the metabolic rate of swimming at low speeds higher than swimming at moderate speeds (Di Santo et al., 2017), but the metabolic rate for hovering in fishes is still under investigation. For bluegill sunfish, Kendall et al. (2007) measured energy consumption at swimming speeds as low as 0.5 L s^-1^. Extrapolating to zero speed, we would estimate that hovering represents about 25% of the total energy consumed. One way a fish may be able to reduce the metabolic cost of hovering is to assume a posture that more closely aligns the COM and COB, thus minimizing destabilizing torques. We wondered, could a neutrally buoyant fish that is unstable in a horizontal posture rest at a pitched equilibrium posture – either stable or unstable equilibrium --to minimize the metabolic costs associated with hovering? We hypothesized that such a fish would minimize the energy spent maintaining a position by resting at stable equilibrium, a posture which is passively maintained. Alternatively, they could rest at an unstable equilibrium. Like a stable equilibrium, torques are zero at an unstable equilibrium, but unlike the stable posture, fish would have to work actively to maintain their posture. The stabilizing torques required near the unstable equilibrium would still be lower than those required to maintain some other posture, and so it might still help to reduce energy cost. To test these hypotheses, we measured the pitch angle of the long axis of the body of 5 bluegill sunfish over the course of 24 hours. Bluegill exhibit a diurnal locomotor activity with fish roughly twice as active during the day than at night (Davis, 1964; Reynolds & Casterlin, 1976). We also measured the approximate position of the COM relative to the COB in the fish to determine the pitch angles correlating with stable and unstable equilibrium for the fish body. Previous studies (Fath et al., 2023) have shown that the COM of bluegill is posterior and ventral to the COB, indicating that stable equilibrium is a head-up posture. We predicted that bluegill would rest with the body in this head-up posture, reducing the horizontal distance between the COM and COB and reducing the amount of torque the fish needs to generate to maintain that posture.

## Methods

We collected 5 bluegill sunfish (*Lepomis macrochirus*) with a seine from the littoral zone of White Pond in Concord, Massachusetts, in the summer of 2019. Prior to testing, the fish were held in separate 10-gallon tanks for at least 2 months and fed five days a week. The water in the tank system was filtered with mechanical and carbon filters. Fish were cared for and handled according to IACUC protocol M2021-99. Fish were maintained on a 12-hour light dark cycle with day running from 7a to 7p and night running from 7p to 7a.

To measure fish body posture over the course of 24 hours we set up a viewing tank (Fig. 1A). The tank water was filtered and aerated. To minimize any flow in the tank due to filtration or aeration, we placed batting, a flow straightener, and thin plastic sheets between the filter and aerator and the filming section of the tank. The tank temperature was 19-20° C. Previous studies found no effect of temperature on diurnal locomotor activity in bluegill (Reynolds & Casterlin, 1976). Fish were fasted for 24 hours prior to the start of the experimental period. The fish was placed into the test tank between noon and 5pm on day zero. The fish was allowed to acclimate to the new tank overnight. Data collection began at 7:00 am, the onset of “day” on day 1. An analog dial on a function generator was used to create a triggering pulse for the cameras at approximately once per minute. Images were collected every 65s ± 7s. The duration and onset of day (7a-7p) and night (7p-7a) were consistent between animal care and experimental conditions. Room lights automatically turned off between 7pm and 7am so that the fish were undisturbed for the entire 24 hours. Three infrared lights (850 nm; CMVision, Houston, TX) were used to illuminate the fish during the 12-hour night period.

**Figure 1.**
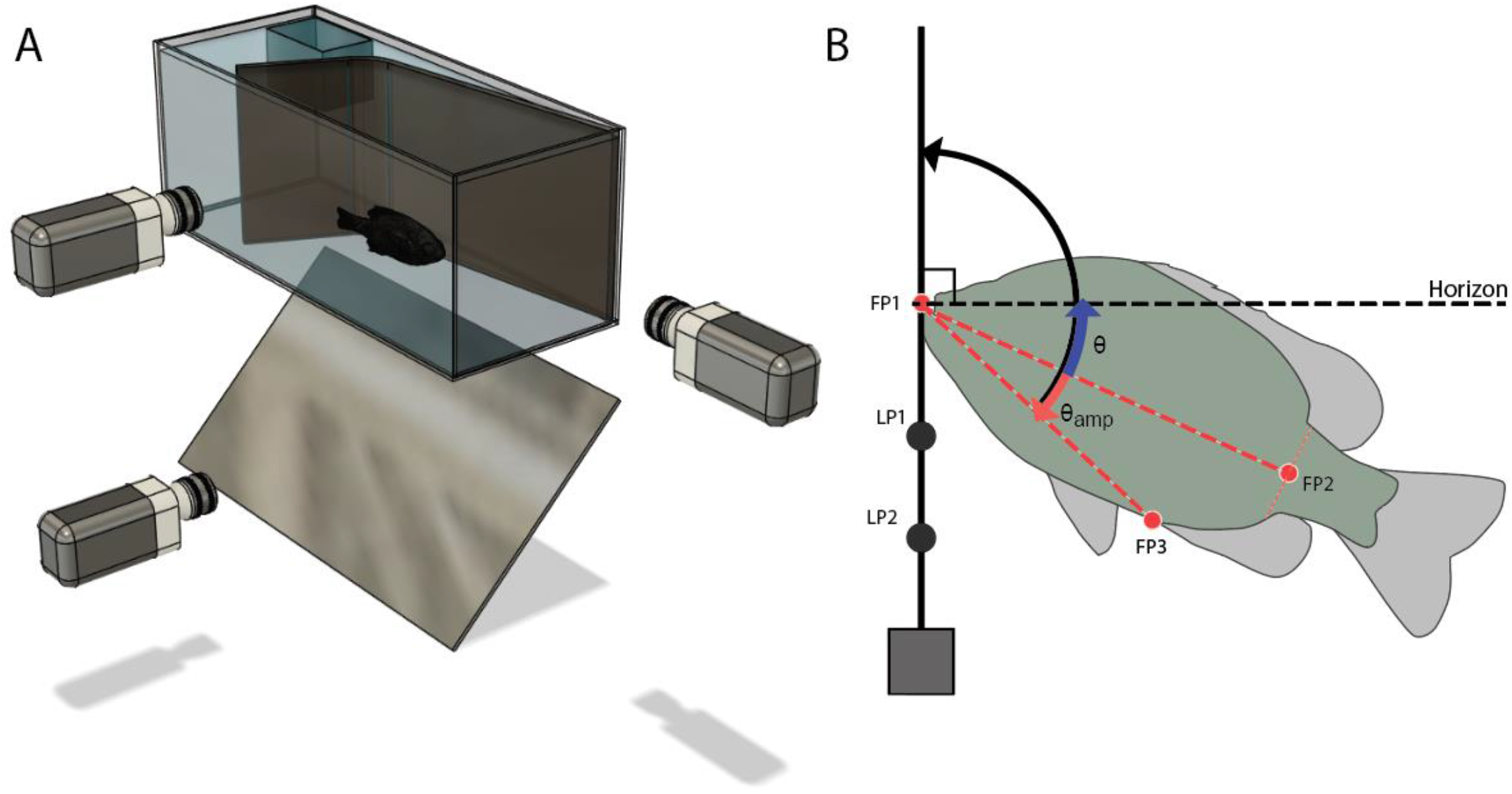
Schematics of (A) the overnight test tanks and (B) the points used to estimate fish pitching angle. A plumb line was suspended in the tank so that it was visible in all three cameras. It was labeled with two points LP1 and LP2, which were used to establish the gravitational vector. Three points were used to calculate the vector of the long axis of the fish body. The tip of the mandible (FP1), a point on the caudal peduncle midway between the posterior base of the anal fin and the posterior base of the dorsal fin (FP2), and the base of the anal fin (FP3).

We imaged the fish with three cameras (Phantom Miro M320 and M120, Vision Research) placed at 90 degrees with respect to each other: aimed at the front, the side, and the bottom of the tank (Fig. 1A). We used DeepLabCut (Mathis et al., 2018) to track three points on the fish body (Fig. 1B): the tip of the mandible (FP1), the mid-peduncle (FP2), and the anal fin base (FP3). These points were used to create a vector following the long axis of the fish body. A weighted plumb line marked with two points (LP1 and LP2) was placed in the view of all three cameras (Fig. 1B). The two points were used to create a vertical gravitational vector.

Using the points established above, we calculated the angle between the gravitational vector and a vector connecting the anal fin base and the anterior-most point of the mandible using the dot product. The long axis of the body was defined as a vector passing from the mandible (FP1) to the mid-peduncle point (FP2). We could not directly measure the midline vector because the mid-peduncle point was not visible from the ventral view and therefore could not be reconstructed in 3D. To get the angle between the long axis of the body and gravitational vector we subtracted the angle from the anal fin to the mandible to the caudal peduncle. Finally, we subtracted 90 to get the angle with the horizon (Equation 1).

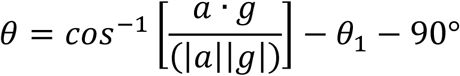

Where a is a vector from the snout of the fish to the base of the anal fin and g is a vector of two points on a plumb line, θ_1_ is the angle FP3-FP1-FP2, and 90° is the angle between the gravitational vector and the horizon.

To estimate the equilibria postures, we transferred the fish to a small tank filled with 7L of tank water with buffered MS-222 (Acros Organics) solution (0.02%) immediately after the 24-hour observational period. After placing the fish in the tank, we waited for all fin movement and gilling to stop. Once the animal was completely immobile, we began to manipulate body posture. Using a large pair of forceps, we gently held the fish at different pitch angles. Then we released the anesthetized fish and measured the change in pitch angle by filming the fish using a DSLR camera filming at 30 fps (Nikon D1500). The camera was set up perpendicular to the sagittal plane of the fish. To measure the moment of release, we placed a mirror under the tank so that we could also see when the forceps were no longer touching the fish body. If the fish rolled instead of pitched, the trial was discarded. We then analyzed the video using FIJI (Schindelin et al., 2012) to quantify the pitch angle of the long axis of the body at the moment of release and after. To measure the direction of pitching it was necessary to wait long enough that the fish was clearly rotating but also soon enough to minimize the influence of other forces (trimming, drag forces) which build up as the fish sinks. We chose 0.33 seconds after release as a time point which balanced those needs. We then plotted the release angle against the change in pitch angle after 0.33 seconds. We fit a regression line to that data, and the point at which the line crossed the X-axis was taken as the equilibrium angle.

### Statistical Analysis

We compared daytime posture to nighttime posture. Comparing day to night, because the angles were not normally distributed, we first used a one sample Kolmogorov-Smirnov test to compare day and night distributions individually to a normal distribution. We then used a Wilcoxon ranked sum test to compare the day angles to the night angles of all 5 fish. Day and night pitching angles are presented as median and standard error of the median (Sokal & Rohlf, 1995).

## Results

### Activity and posture vary between day and night

We measured the activity and posture of 5 bluegill with mean mass of 55 ± 41g (mean ± standard deviation) and a mean size of 11 ± 3 cm over the course of 24 hours. We used displacement from one image to the next (taken at 1 frame per minute) as a measure of activity. We found that fish were more active during the day than at night, moving 0.46 ± 0.19 BL/min (median ± standard error of the median) (Fig. 2A). At night, fish were less active, moving 0.33 ± 0.21 BL/min on average, although there were peaks in activity (around 18hours in Fig.2A).

**Figure 2.**
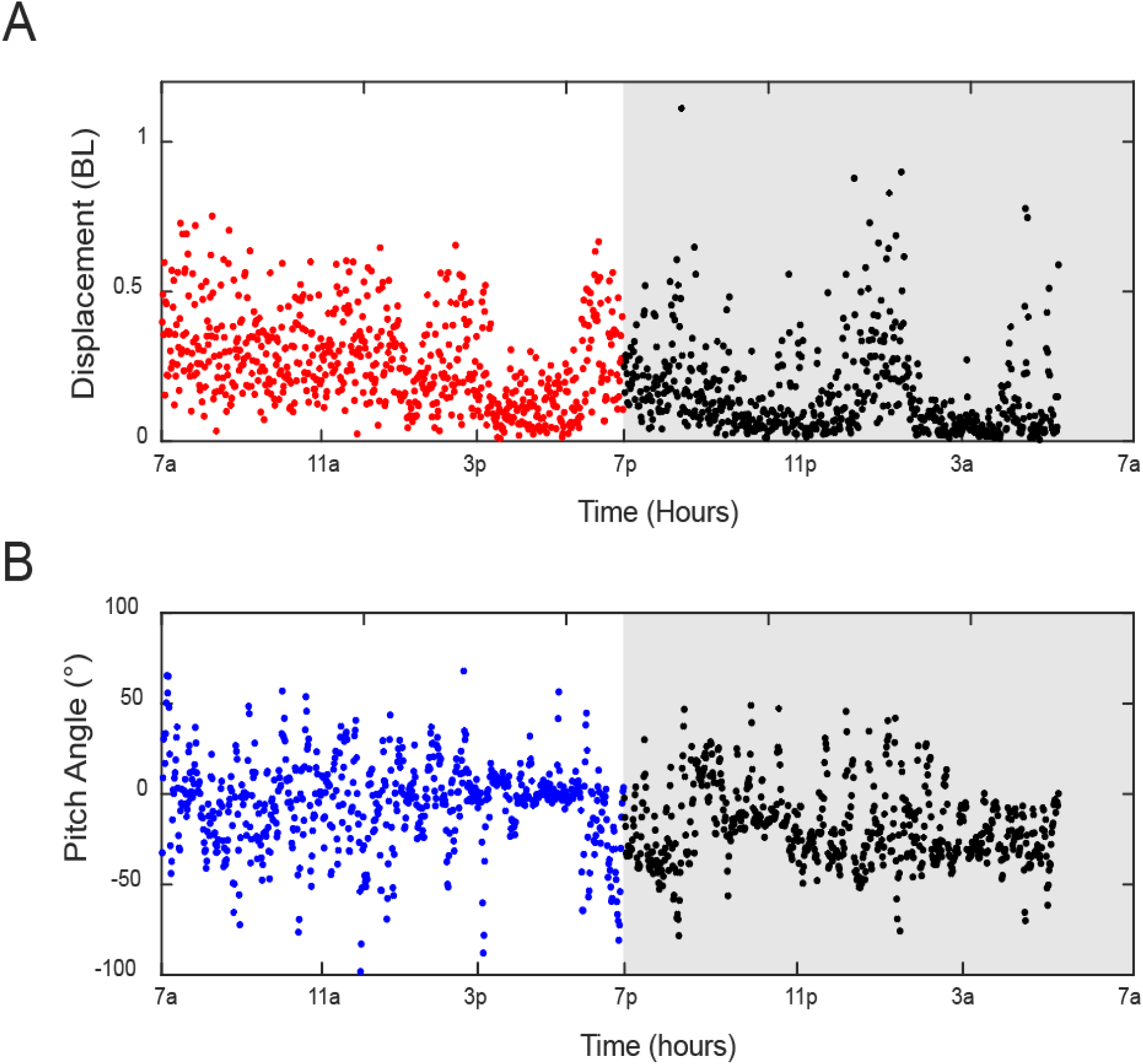
Displacement (A) and pitch angle (B) of one bluegill over the course of 24 hours. Data collected during the day (7a to 7p) are indicated by colored points on a white background. Data collected at night (between 7p and 7a) are indicated by black points on a gray background. z

Bluegill had significantly different postures at night compared to daytime postures. We found that the day and night pitch angles have a non-normal distribution (one sample Kolmogorov Smirnov test; p < 0.001). The fish maintained a head-down pitch angle on average at all times: -3.4±0.8° (negative angles indicate head down) during the day and -10.7± 0.4° at night (Fig. 2B). Four of five fish were pitched significantly further head down at night (Wilcoxon Ranked Sum Test, p = <0.001), with one fish with a pitch angle that was not significantly different between day and night (p=0.37).

### Measuring stable and unstable equilibrium angles

We also measured the pitch angle of stable and unstable equilibria for anesthetized bluegill (Fig. 3). We held each fish at different pitch angles, released it, and recorded the change in pitch angle after 0.33 s. All fish were negatively buoyant and sank slowly upon release. Stable equilibrium was found to be a ventral side up (belly up) posture. The unstable equilibrium angle was estimated to be -53±26°, a ventral-side-down posture with the head pointed down.

**Figure 3.**
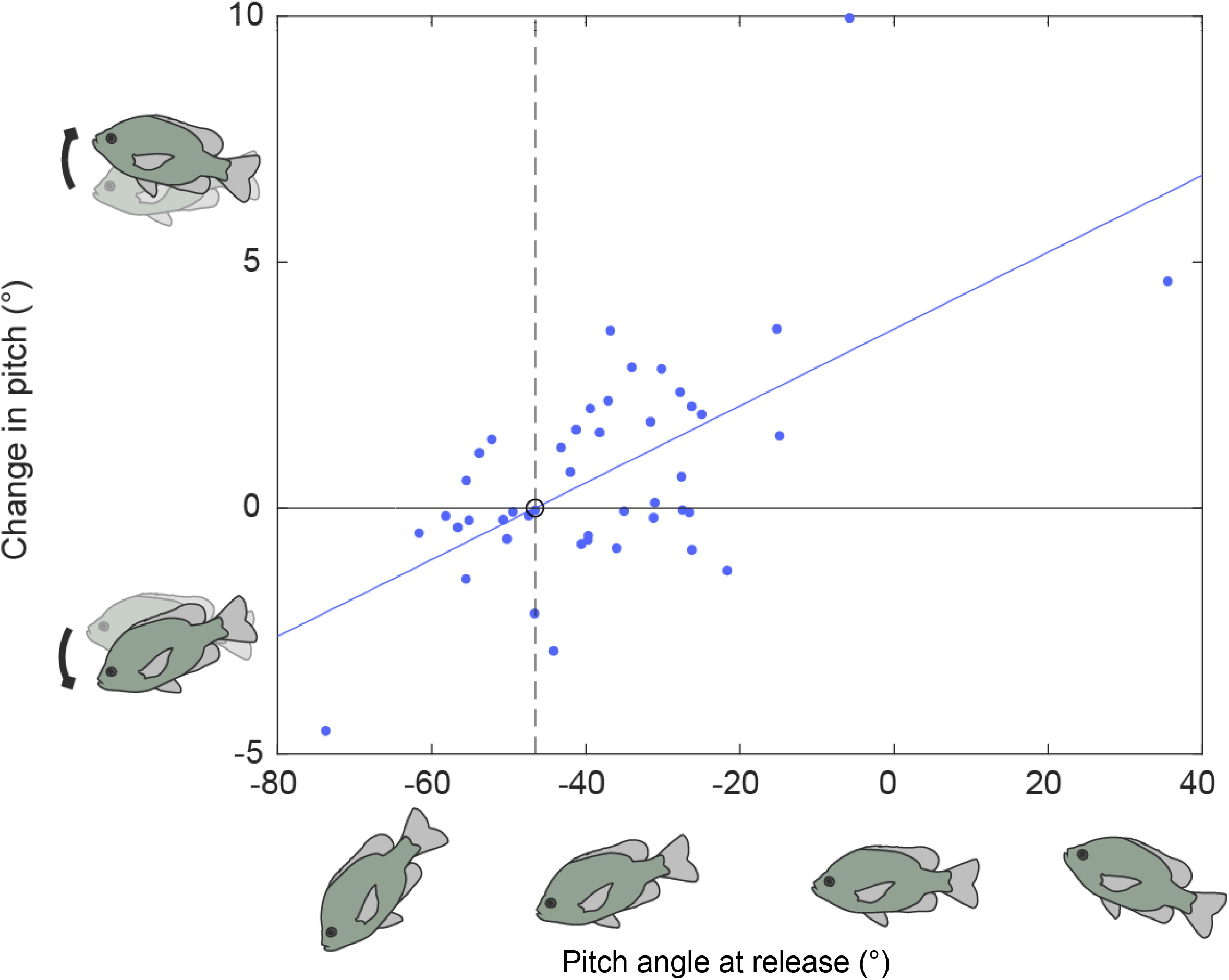
The change in pitch angle of a bluegill when released in a static tank with a pair of forceps. The X-intercept of the regression indicates the release angle which is estimated to be unstable equilibrium. Positive values on the Y-axis indicate a head-up change in pitch and negative values indicate a head-down change pitch angle. The change in pitch angle was measured 0.33 s after release.

Finally, we compared the pitch angles of the body during the day and night with the unstable equilibrium angle as measured when anesthetized (Fig. 4). The average body angle during the day was the highest, -3.4±0.8° pitched head-down; the body angle during the night was significantly lower, at -10.7± 0.4°; but the angle of the anesthetized fish at its unstable equilibrium was lower, at -53±26° pitched head-down. We compared the unstable equilibrium angle, estimated from the regressions for each fish (example in Fig. 3), to the distribution of posture angles taken by each fish at night. We found that the unstable equilibrium angle was steeper than 93% of all postures taken by one fish, and steeper than 97% of the postures taken at night in the 4 other fish.

**Figure 4.**
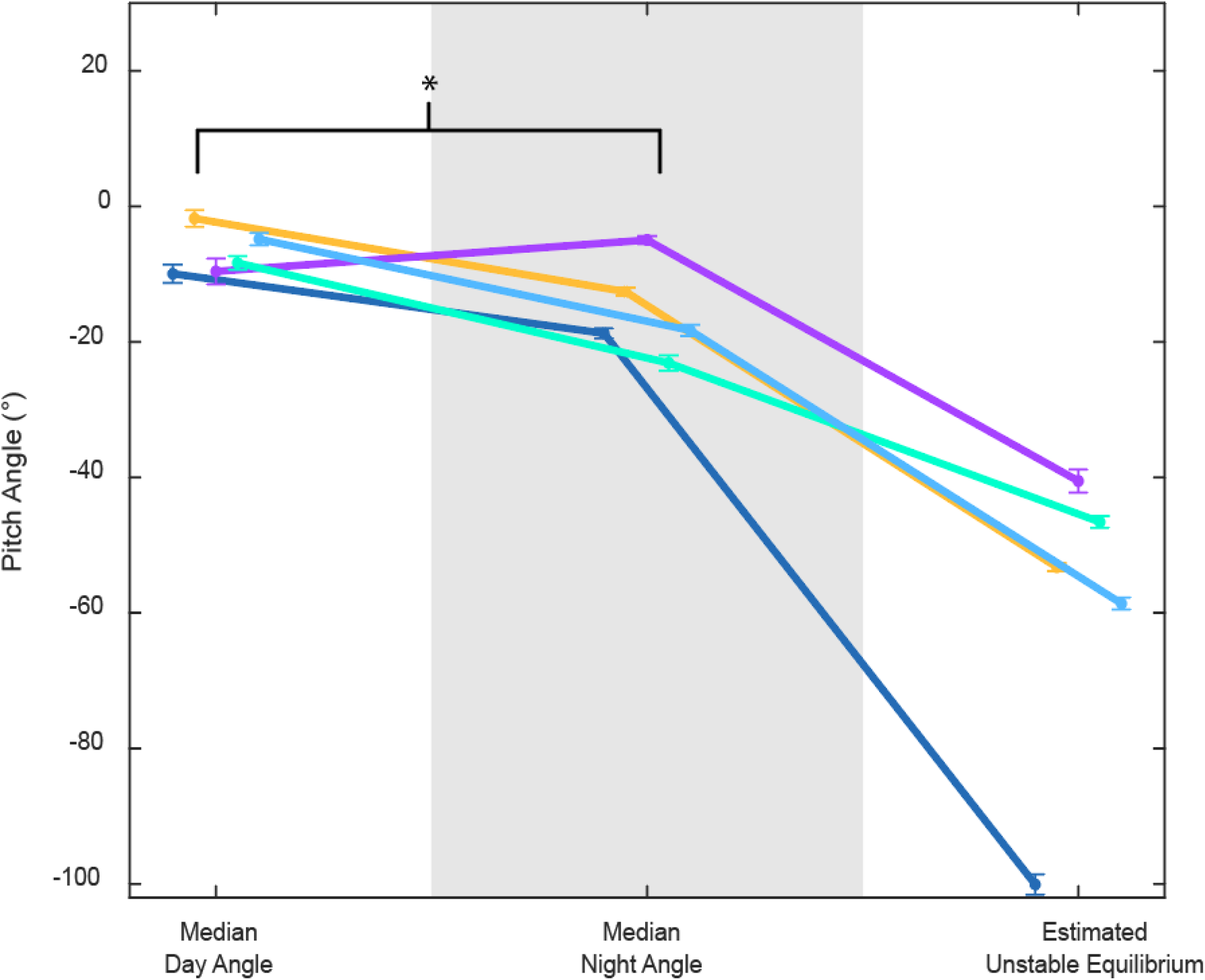
The pitch angle for 5 bluegill during the day, night, and at unstable equilibrium. The day and night angle values are the median values as calculated from the postures collected during the day and night periods for each fish and the error bars are the standard error of the median. The values for the unstable equilibrium posture are calculated from the linear regression the point at which there is no change in posture. The error bars are the SEM.

## Discussion

We found that at night bluegill rest pitched at -10.7± 0.4° (where negative angles indicate a head down posture). This is pitched significantly more head-down than the daytime angle of -3.4± 0.8°. Contrary to our hypothesis, this posture is not equal to either their stable or unstable equilibrium postures. The stable equilibrium in bluegill is a belly-up posture. We never observed unanesthetized bluegill in a belly up posture and previous studies indicate that bluegill (and other fishes) are highly motivated to avoid rolling (Eidietis et al., 2002; Webb, 2004).

Many terrestrial animals and some fishes sleep in passive postures (Kelly et al., 2022; Shapiro & Hepburn, 1976), however bluegill rest at an unstable posture which requires active maintenance. This suggests that there may be a benefit to keeping a specific posture which outweighs the metabolic cost of maintaining that unstable posture. The unstable state confers the ability to rapidly change that state, making the fish more maneuverable and more difficult for a predator to catch (Clemente & Wilson, 2016). Bluegill share habitats with piscivorous nocturnal predators (Keast et al., 1987) and fish can be more vulnerable to predation in the evening and night early night (Holbrook & Schmitt, 2002). Maintaining a more maneuverable state even while resting may let the bluegill keep an escape advantage over predators (Howland 1974).

While bluegill do tend to rest at more steeply pitched angles at night, they do not rest at the unstable equilibrium, which was -53±26° on average. Resting at an unstable equilibrium would mean that there are no net torques on the body and would allow the bluegill to maintain maneuverability while also minimizing the metabolic of hovering. However, this is not what they do. They rest at -10.7± 0.4°, an angle that is closer to the unstable equilibrium than the daytime posture, but still substantially different from the equilibrium.

There are a few possible explanations for this discrepancy. First, there may be a difference in equilibrium posture of live and anesthetized fish. Postural equilibrium is the position at which all forces are balanced, generating no torques. Our anesthetized pitching experiments allow us to able to directly observe the balance of gravity and buoyancy on a fish body. However, there are mechanical differences between an active and an anesthetized fish which may alter the balance of forces on the body. Bluegill are near neutrally buoyant but ultimately are negatively buoyant fish with a density of 1.0016 ± 0.0015 g/cm^3^ (Webb, 1993), so fish must generate lift as they rest. Additionally, even small flows such as those involved in respiration (Tytell & Alexander, 2007) have the potential to interact with the fins and body, slightly changing the balance of forces on the body. Lift generation and gilling are two examples of forces which, added to the gravitational and buoyant forces, may change equilibrium posture of the active fish compared to the anesthetized fish.

Alternately, if we assume that the unstable equilibrium posture measured for the anesthetized fish is representative of the unstable equilibrium posture of the live fish, this may indicate that the activity needed to maintain that posture is metabolically negligible. In bluegill as in many fishes the COM and COB are close to one another, so they generate relatively low torques even in the most unstable postures (Fath et al., 2023). However, the relationship between metabolism and posture and how those change during activity and rest in fishes must be directly examined. Bluegill resting metabolism has been studied (Jones et al., 2007; Kendall et al., 2007), but not specifically while the fish is hovering. Due to the sporadic nature of fish movement, activity is often not quantified while recording resting metabolism (Chabot et al., 2016). Extrapolating from bluegill swimming data, the total metabolic power needed to swim at 0 swimming speed is estimated to be 0.47 W kg^-1^ (Kendall et al., 2007), but it is not clear how well this would reflect the actual metabolic energy demands of hovering. Additionally, extrapolating resting metabolic data from swimming data may (Roche et al., 2013; Schurmann & Steffensen, 1997) or may not (Duthie, 1982; Roche et al., 2013) agree with measurements made at zero swimming speed. Fishes generally have a lower metabolism at night (Grantner & Taborsky, 1998; Kelly et al., 2022), but the role posture may play, if any, is not known. Measuring oxygen consumption (MO_2_) while also recording posture and kinematics over the course of 24 hours would provide insight into the differing metabolic costs at various behaviors at or near zero swimming speed during periods of rest and activity.

The equilibrium postures seen in the anesthetized pitching experiments indicate that the bluegill’s COM is posterior and dorsal to the COB, a pattern somewhat different from what was estimated in both previous studies (Webb and Weihs, 1994; Fath et al., 2023) of their locations in bluegill sunfish. This configuration was consistent across all 5 fish measured. Using the plumb line method, Webb and Weihs (1994) found no statistical difference between the location of the COM and COB in bluegill, suggesting that bluegill may be neutrally stable. Using µCT reconstructions, Fath et al. (2023) found that the COM was significantly posterior and ventral to the COB in most bluegill. This would result in an unstable equilibrium posture with the fish pitched steeply head-up. Instead, our anesthetized pitching experiments found that bluegill had an unstable equilibrium posture pitched steeply head-down, indicating that the COM is posterior and dorsal to the COB in all the fish we measured, rather than posterior and ventral as estimated by Fath et al. (2023). The anesthetized pitching experiments used here cannot tell us the exact locations or distances between the COM and COB. However, the current measurements *in situ* may more accurately represent the relative locations of the COM and COB. The plumb line and CT methods use dead fish measured in air to calculate COM and COB location and estimate static equilibrium postures. The swim bladder is mobile in the body cavity (Fath et al., 2023), and very small changes between live animals in situ and dead animals used for scanning may affect the estimated locations and postures. Future researchers should consider their µCT protocol carefully, and should ideally scan live fish in water, at a horizontal orientation.

## Conclusion

Bluegill rest at a significantly more pitched head down angle at night compared to their median posture during the day, with less variability in posture. This does not vertically align the COM and COB, but may move the points closer together compared to day-time postures, lowering the energy needed to create stabilizing torques.

## References

Chabot, D., Steffensen, J. F., & Farrell, A. P. (2016). The determination of standard metabolic rate in fishes. Journal of Fish Biology, 88(1), 81–121. https://doi.org/10.1111/jfb.12845

Ciancio, J. E., Venerus, L. A., Trobbiani, G. A., Beltramino, L. E., Gleiss, A. C., Wright, S., Norman, B., Holton, M., & Wilson, R. P. (2016). Extreme roll angles in Argentine sea bass: Could refuge ease posture and buoyancy control of marine coastal fishes? Marine Biology, 163(4), 90. https://doi.org/10.1007/s00227-016-2869-z

Clark, C. J., & Dudley, R. (2010). Hovering and forward flight energetics in Anna’s and Allen’s hummingbirds. Physiological and Biochemical Zoology, 83(4), 654–662. https://doi.org/10.1086/653477

Clemente, C. J., & Wilson, R. S. (2016). Speed and maneuverability jointly determine escape success: Exploring the functional bases of escape performance using simulated games. Behavioral Ecology, 27(1), 45–54. https://doi.org/10.1093/beheco/arv080

Davis, R. E. (1964). Daily “Predawn” Peak of Locomotion in Fish. Animal Behavior, 12(2–3), 272–283.

Di Santo, V., Kenaley, C. P., & Lauder, G. V. (2017). High postural costs and anaerobic metabolism during swimming support the hypothesis of a U-shaped metabolism–speed curve in fishes. Proceedings of the National Academy of Sciences of the United States of America, 114(49), 13048–13053. https://doi.org/10.1073/pnas.1715141114

Dorfman, A., Hills, T. T., & Scharf, I. (2022). A guide to area-restricted search: a foundational foraging behaviour. Biological Reviews, 97(6), 2076–2089. https://doi.org/10.1111/brv.12883

Duthie, G. G. (1982). The Respiratory Metabolism of Temperature-Adapted Flatfish at Rest and During Swimming Activity and the Use of Anaerobic Metabolism at Moderate Swimming Speeds. In J. exp. Biol (Vol. 97).

Eidietis, L., Forrester, T. L., & Webb, P. W. (2002). Relative abilities to correct rolling disturbances of three morphologically different fish. Canadian Journal of Zoology, 80(12), 2156–2163. https://doi.org/10.1139/z02-203

Fath, M. A., Nguyen, S. V, Donahue, J., McMenamin, S. K., & Tytell, E. D. (2023). Static Stability and Swim Bladder Volume in the Bluegill Sunfish (Lepomis macrochirus). Integrative Organismal Biology, 5(1). https://doi.org/10.1093/iob/obad005

Fish, F. E., & Holzman, R. (2019). Swimming turned on its head: Stability and maneuverability of the shrimpfish (Aeoliscus punctulatus). Integrative Organismal Biology, 1(1). https://doi.org/10.1093/iob/obz025

Goldshmid, R., Holzman, R., Weihs, D., & Genin, A. (2004). Aeration of corals by sleep-swimming fish. Limnology and Oceanography, 49(5), 1832–1839. https://doi.org/10.4319/lo.2004.49.5.1832

Grantner, A., & Taborsky, M. (1998). The metabolic rates associated with resting, and with the performance of agonistic, submissive and digging behaviours in the cichlid Fish Neolamprologus pulcher (Pisces: Cichlidae). Journal of Comparative Physiology, 168, 427–433.

Holbrook, S. J., & Schmitt, R. J. (2002). Competition for shelter space causes density-dependent predation mortality in damselfishes. Ecology, 83(10), 2855–2868. https://doi.org/10.1890/0012-9658(2002)083[2855:CFSSCD]2.0.CO;2

Howland, H. C. (1974). Optimal Strategies for Predator Avoidance : The Relative Importance of Speed and Manoeuvrability. In J. theor. Biol (Vol. 47).

Joiner, W. J. (2016). Unraveling the Evolutionary Determinants of Sleep. In Current Biology (Vol. 26, Issue 20, pp. R1073–R1087). Cell Press. https://doi.org/10.1016/j.cub.2016.08.068

Jones, E. A., Lucey, K. S., & Ellerby, D. J. (2007). Efficiency of labriform swimming in the bluegill sunfish (Lepomis macrochirus). Journal of Experimental Biology, 210(19), 3422–3429. https://doi.org/10.1242/jeb.005744

Keast, A., Harker, J., & Turnbull, D. (1987). Nearshore fish habitat utilization and species associations in Lake Opinicon (Ontario, Canada). Environmental Biology of Fishes, 3(2), 173–184.

Keene, A. C., & Duboue, E. R. (2018). The origins and evolution of sleep. In Journal of Experimental Biology (Vol. 221, Issue 11). Company of Biologists Ltd. https://doi.org/10.1242/JEB.159533

Kelly, M. L., Collin, S. P., Hemmi, J. M., & Lesku, J. A. (2020). Evidence for Sleep in Sharks and Rays: Behavioural, Physiological, and Evolutionary Considerations. In Brain, Behavior and Evolution (Vol. 94, Issues 1–4, pp. 37–50). S. Karger AG. https://doi.org/10.1159/000504123

Kelly, M. L., Collins, S. P., Lesku, J. A., Hemmi, J. M., Collin, S. P., & Radford, C. A. (2022). Energy conservation characterizes sleep in sharks. Biology Letters, 18(3). https://doi.org/10.1098/rsbl.2021.0259

Kendall, J. L., Lucey, K. S., Jones, E. A., Wang, J., & Ellerby, D. J. (2007). Mechanical and energetic factors underlying gait transitions in bluegill sunfish (Lepomis macrochirus). Journal of Experimental Biology, 210(24), 4265–4271. https://doi.org/10.1242/jeb.009498

Mathis, A., Mamidanna, P., Cury, K. M., Abe, T., Murthy, V. N., Mathis, M. W., & Bethge, M. (2018). DeepLabCut: markerless pose estimation of user-defined body parts with deep learning. Nature Neuroscience, 21(9), 1281–1289. https://doi.org/10.1038/s41593-018-0209-y

Nathan, R., Getz, W. M., Revilla, E., Holyoak, M., Kadmon, R., Saltz, D., & Smouse, P. E. (2008). A movement ecology paradigm for unifying organismal movement research. http://www.pnas.orgcgidoi10.1073pnas.0800375105

Pyke, G. H. (1983). Animal movements - an optimal foraging theory approach. In The ecology of animal movement (pp. 7–31). Elsevier. https://doi.org/10.1016/B978-0-12-809633-8.90160-2

Reynolds, W. W., & Casterlin, M. E. (1976). Locomotor Activity Rhythms in the Bluegill Sunfish, Lepomis macrochirus. Source: The American Midland Naturalist, 96(1), 221–225.

Roche, D. G., Binning, S. A., Bosiger, Y., Johansen, J. L., & Rummer, J. L. (2013). Finding the best estimates of metabolic rates in a coral reef fish. Journal of Experimental Biology, 216(11), 2103–2110. https://doi.org/10.1242/jeb.082925

Schindelin, J., Arganda-Carreras, I., Frise, E., Kaynig, V., Longair, M., Pietzsch, T., Preibisch, S., Rueden, C., Saalfeld, S., Schmid, B., Tinevez, J. Y., White, D. J., Hartenstein, V., Eliceiri, K., Tomancak, P., & Cardona, A. (2012). Fiji: An open-source platform for biological-image analysis. In Nature Methods (Vol. 9, Issue 7, pp. 676–682). https://doi.org/10.1038/nmeth.2019

Schmidt, M. H. (2014). The energy allocation function of sleep: A unifying theory of sleep, torpor, and continuous wakefulness. In Neuroscience and Biobehavioral Reviews (Vol. 47, pp. 122–153). Elsevier Ltd. https://doi.org/10.1016/j.neubiorev.2014.08.001

Schurmann, H., & Steffensen, J. F. (1997). Effects of temperature, hypoxia and activity on the metabolism of juvenile Atlantic cod. Journal of Fish Biology, 50(6), 1166–1180. https://doi.org/10.1111/j.1095-8649.1997.tb01645.x

Shapiro, C. M., & Hepburn, H. R. (1976). Sleep in a Schooling Fish, Tilapia mossambica. In Physiology &Behavior (Vol. 16). Pergamon Press and Brain Research Publ.

Siegel, J. (2011). Sleep in Animals: A State of Adaptive Inactivity. In M. H. Kryger, T. Roth, &W. C. Dement (Eds.), Principles and practice of sleep medicine (5th ed., pp. 126–138). Elsevier Saunders.

Sokal, R. R., & Rohlf, R. J. (1995). Biometry: The Principles and Practice of Statistics in Biological Research (3rd ed.). W.H. Freeman and Company.

Tytell, E. D., & Alexander, J. K. (2007). Bluegill Lepomis macrochirus synchronize pectoral fin motion and opercular pumping. Journal of Fish Biology, 70(4), 1268–1279. https://doi.org/10.1111/j.1095-8649.2007.01416.x

Webb, P. W. (1993). Is tilting behaviour at low swimming speeds unique to negatively buoyant fish? Observations on steelhead trout, Oncorhynchus mykiss, and bluegill, Lepomis macrochirus. Journal of Fish Biology, 43, 687–694.

Webb, P. W. (2004). Response latencies to postural disturbances in three species of teleostean fishes. The Journal of Experimental Biology, 207(1), 955–961. https://doi.org/10.1242/jeb.00854

Webb, P. W., & Weihs, D. (1994). Hydrostatic stability of fish with swim bladders: not all fish are unstable. Canadian Journal of Zoology, 72(6), 1149–1154. https://doi.org/10.1139/z94-153

Weihs, D. (1993). Stability of Aquatic Animal Locomotion.

Zhdanova, I. v, Wang, S. Y., Leclair, O. U., & Danilova, N. P. (2001). Melatonin promotes sleep-like state in zebrafish. In Brain Research (Vol. 903). http://www.bres-interactive.com

